# Sample overlap in genome-wide association meta-analysis: practical comparisons across significance thresholds

**DOI:** 10.64898/2026.10.14.759234

**Authors:** Zikai Xu, Lin Zhang, Lloyd T. Elliott

## Abstract

Meta-analysis is widely used in genome-wide association studies (GWAS) to combine evidence across studies without requiring individual-level data. However, as both the number of studies included in meta-analyses and their sample sizes continue to grow, the same participant may be included in more than one study. Such sample overlap induces dependence across studies and violates the independence assumptions underlying standard meta-analysis methods. HostSeq is a Canadian initiative in which 15 clinical and epidemiological studies were assembled with DNA samples from approximately 10,000 Canadians infected by the SARS-CoV-2 virus. We observed duplicate samples across different participating studies. This observation raises an important practical question: what is the impact of sample overlap on association results, and when should such overlap be a concern? Motivated by this real problem, we sought to compare five meta-analysis methods and evaluate their performance against a reference mega-analysis in which the true sample overlap is known. We show that when the degree of sample overlap is relatively small, its impact on association results is minimal, even at the genome-wide significance threshold of *α* = 5 × 10^−8^. We demonstrate these findings through extensive simulation studies and an application to overlapping cohorts within HostSeq.

**Author summary:** Combining results from multiple genetic studies can help researchers identify genetic variants associated with diseases without requiring access to individual participants’ data. However, some individuals may be involved in more than one study, creating dependence between study results. Ignoring this dependence may lead to misleading conclusions about genetic associations. Motivated by overlapping participants identified in the Canadian HostSeq initiative, we investigated how sample overlap affects genetic association results and compared existing statistical methods that account for it. Using simulations and HostSeq genetic data, we found that small sample overlaps may have little impact on association results, but ignoring larger overlaps can lead to reporting associations when none exist. This problem can become more pronounced under the stringent significance thresholds commonly used in genetic studies. We also found that existing methods for correcting sample overlap have different strengths and limitations. Our findings provide practical guidance on when sample overlap may require correction and which statistical methods researchers should consider.

## Introduction

Genome-wide association studies (GWAS) have become a central tool for investigating the genetic basis of complex traits and diseases. Meta-analysis combines evidence from **multiple studies without requiring individual-level data to be shared [1]. In practice**, sample overlap can occur in genetic studies due to the repeated inclusion of participants across multiple cohorts, biobanks, or research initiatives. HostSeq is a Canadian initiative that assembled whole-genome sequencing and clinical data from approximately 10,000 participants recruited through multiple COVID-19 studies [2, 3]. In the HostSeq databank, some participants were recruited into more than one contributing study. For example, the Quebec COVID-19 Biobank (BQC19) and the IRCM Post-COVID-19 Research Clinic (IPCO) shared 11 participants. Such overlap creates dependence between study results. Although overlapping participants were identified and removed in the original HostSeq analysis, this may not always be feasible in other meta-analyses. Motivated by this practical challenge, we investigate how the magnitude of sample overlap affects meta-analysis results and when adjustment for such overlap becomes necessary.

Researchers have considered statistical adjustment to account for dependence induced by overlapping participants. Lin and Sullivan [4] introduced an adjusted meta-analysis approach designed specifically for case-control studies with partially overlapping participants. This method combines the estimated log-odds ratios from each study and accounts for their induced correlation due to shared participants. It combines the effect estimates using covariance-adjusted inverse-variance weighting, followed by a Wald test for inference. However, the overlap structure required for this adjustment, including the separate counts of shared cases and controls, may not be available, limiting the method’s applicability.

Combining study-level *p*-values provides another approach to meta-analysis through global hypothesis testing. Each input *p*-value measures evidence against a study-specific null hypothesis. A combination method summarizes these values in a test statistic and produces an overall *p*-value for the global null hypothesis. In our setting, this global null states that the variant is not associated with the trait in any of the studies; rejection indicates evidence of association in at least one study. Many methods have been developed for combining independent tests of significance [5–9], including weighted combinations [10–12]. Their relative performance depends on the pattern of associations across studies [13, 14]. We focus on Fisher’s method, a classical and widely used approach that combines *p*-values through the sum of their negative logarithms.

Sample overlap induces dependence among study results, violating the independence assumption of standard *p*-value combination methods. Ignoring this dependence can lead to inflated false-positive rates. Several extensions account for correlated *p*-values, including Hartung’s inverse-normal approach [15] and Brown’s adaptation of Fisher’s method [16]. Brown approximated the distribution of Fisher’s statistic under the global null using a rescaled chi-square distribution. Subsequent work developed approximations for the covariance between the −2 log *p*_*i*_ terms [17] and an empirical adaptation of Brown’s method [18]. These developments allow dependence to be incorporated, although the accuracy of the combined *p*-values remains a consideration when choosing an approximation [19]. Brown’s approximation can be inaccurate in the tail, leading to inflated T1E at stringent significance thresholds such as those used in GWAS. To address this limitation, Zhang and Wu [20] proposed moment-ratio (MR) and hybrid (HYB) methods for approximating the null distribution of the combined test statistic within the generalized Fisher (GFisher) framework, and compared them with the generalized Brown (GB) approximation. These methods accommodate correlated *p*-values in general, rather than sample overlap specifically. Their application to overlapping-study meta-analysis requires an estimate of the correlation among the study-level test statistics.

In this study, we investigate how overlapping participants affect GWAS meta-analysis and evaluate existing methods for accounting for sample dependence. We first review the statistical formulations of Fisher’s method, three GFisher-based methods (GB, HYB, and MR), and Lin and Sullivan’s adjusted-meta. Assuming that overlapping participants are included twice at random, we approximate the correlation between study-specific score test statistics using the number of overlapping participants and the individual study sizes. We then conduct simulation studies to evaluate type I error (T1E) and power across different degrees of sample overlap and allele frequencies, using mega-analysis based on unique participants as the benchmark. We further evaluate these methods using HostSeq genetic data and cohort sample sizes, with additional GWAS-sized simulations to evaluate the accuracy of these methods at stringent significance levels. Finally, we discuss practical recommendations for handling sample overlap in GWAS meta-analysis, the limitations of our analyses, and directions for future research.

## Materials and methods

### Problem setup and notations

We here consider a binary phenotype outcome. The two overlapping studies, 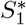 and 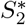, with sample sizes 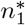 and 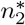, respectively, share *n*_12_ participants. We denote the shared participants as *S*_12_, while 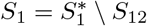is the sample that is unique to Study 1 and 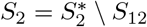 is the sample that is unique to Study 2. Their union contains 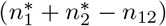 distinct participants (see Table 1 for the notations). Here we assume that any participant in either study has an equal chance to be included in the overlapping sample, *S*_12_.

**Table 1.**
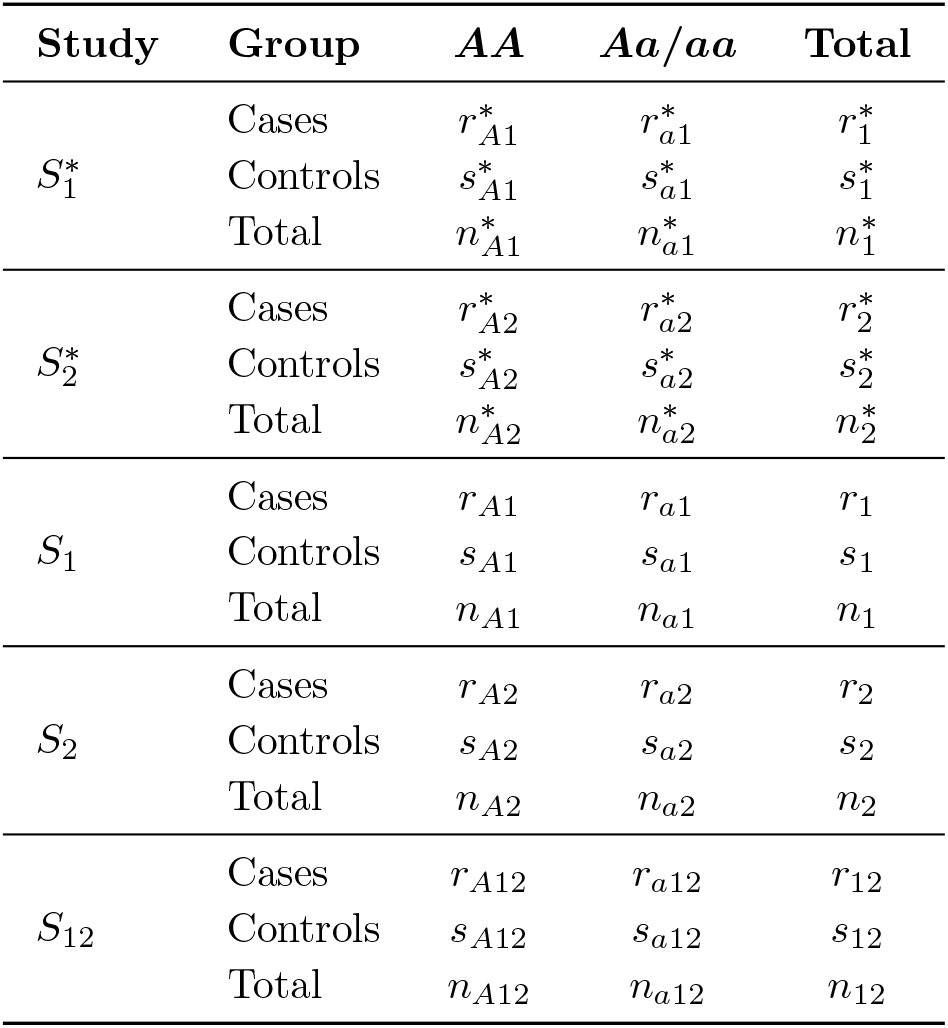
Genotype count notations in two partially overlapping case-control studies. The observed studies are denoted by 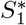 and 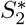, their shared participants by *S*_12_, and their study-specific components by 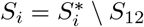 for *i* = 1, 2. The minor allele is denoted by *A*. Subscripts *A* and *a* identify the *AA* and combined *Aa/aa* genotype groups, respectively. Symbols *r, s*, and *n* denote case, control, and total counts. Corresponding counts satisfy 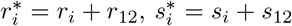, and 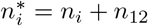

| Study | Group | $AA$ | $Aa/aa$ | Total |
| --- | --- | --- | --- | --- |
| $S_1^*$ | Cases | $r_{A1}^*$ | $r_{a1}^*$ | $r_1^*$ |
| | Controls | $s_{A1}^*$ | $s_{a1}^*$ | $s_1^*$ |
| | Total | $n_{A1}^*$ | $n_{a1}^*$ | $n_1^*$ |
| $S_2^*$ | Cases | $r_{A2}^*$ | $r_{a2}^*$ | $r_2^*$ |
| | Controls | $s_{A2}^*$ | $s_{a2}^*$ | $s_2^*$ |
| | Total | $n_{A2}^*$ | $n_{a2}^*$ | $n_2^*$ |
| $S_1$ | Cases | $r_{A1}$ | $r_{a1}$ | $r_1$ |
| | Controls | $s_{A1}$ | $s_{a1}$ | $s_1$ |
| | Total | $n_{A1}$ | $n_{a1}$ | $n_1$ |
| $S_2$ | Cases | $r_{A2}$ | $r_{a2}$ | $r_2$ |
| | Controls | $s_{A2}$ | $s_{a2}$ | $s_2$ |
| | Total | $n_{A2}$ | $n_{a2}$ | $n_2$ |
| $S_{12}$ | Cases | $r_{A12}$ | $r_{a12}$ | $r_{12}$ |
| | Controls | $s_{A12}$ | $s_{a12}$ | $s_{12}$ |
| | Total | $n_{A12}$ | $n_{a12}$ | $n_{12}$ |

We assume a recessive disease model here. However, this can be easily generalized to any disease model such as the additive model. To test the association between the genotype and binary phenotype in Study *i* (for *i* = 1 and 2), the score statistic is

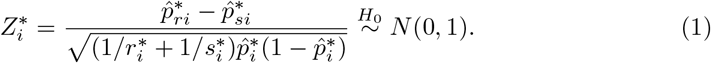

Here,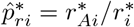and 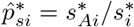 are the frequencies of genotype *AA* among cases and controls in Study *i*, and 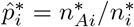 is the pooled frequency. We assume Hardy–Weinberg equilibrium holds in all samples. Under the null hypothesis of no association, 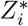 is asymptotically standard normal with *p*-value 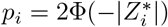, where Φ is the cumulative distribution function (CDF) for *N* (0, 1).

If we partition all the participants in Study 1 and 2 into three independent studies *S*_1_, *S*_2_ and *S*_12_, we can compute their score test statistics *Z*_1_, *Z*_2_ and *Z*_12_ in a similar fashion to (1).

We further show that

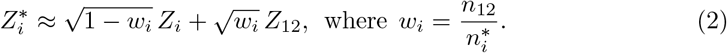

Thus, we can approximate the correlation between the score test statistics of Study 1 and Study 2 by

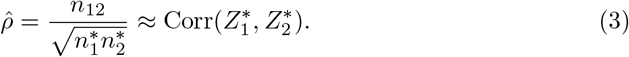

See Section A in S1 File for the derivations.

### Fisher’s combination test and GFisher approximations

In this study, we consider the classical Fisher’s approach to combine the *p*-values from two overlapping studies. For Study *i* = 1, 2, let

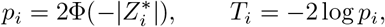

where 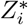 is the score statistic defined in Eq (1) and Φ is the standard normal CDF. The classical Fisher’s test statistic is

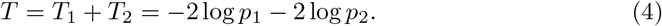

When Study 1 and Study 2 are two independent studies, 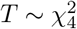 under the null hypothesis of no association, and the corresponding *p*-value is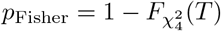.

When Study 1 and Study 2 share overlapping participants, the joint null distribution of the study-level statistics is

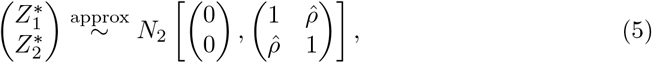

where 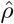 is estimated as in Eq (3). Under the joint distribution of 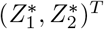 defined in Eq (5), the Fisher’s test statistic *T* has

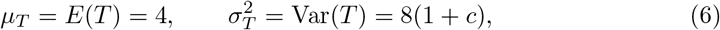

where *c* = Corr(*T*_1_, *T*_2_) is the correlation between the transformed *p*-values.

To calculate *c*, we apply Theorem 1 and Corollary 1 of Zhang and Wu [20]. For our two-study Fisher statistic,

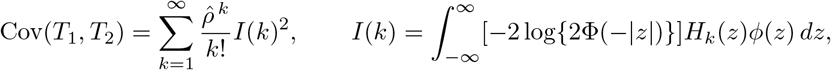

where *ϕ* is the standard normal density and *H*_*k*_ is the probabilists’ Hermite polynomial of order *k*. Only even-order terms contribute because the input *p*-values are two-sided.

GFisher-based methods approximate the null distribution of *T* using Gamma distributions. The GFisher-based *p*-value is defined

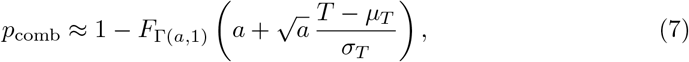

where *F*_Γ(*a*,1)_ is the CDF of Γ(*a*, 1). We next introduce three GFisher-based approaches with different ways to approximate the shape parameter *a*.

### Generalized Brown’s (GB) method

The original Brown’s method approximates the null distribution of Fisher’s statistic for dependent *p*-values using a scaled *χ*^2^ distribution [16], which the test statistic *T* in Eq (4) is approximated by 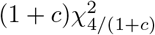 . Generalized Brown’s method extends Brown’s mean-variance matching approach to a broader class of statistics. In our case of combining two overlapping association studies, GB and the original Brown’s method adopt the same distribution to match the mean and variance of *T* [16, 20]. *The corresponding shape parameter is*

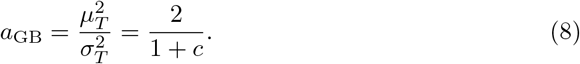

In our analysis, we implemented GB using the R package GFisher, version 0.2.0.

### Moment-ratio matching (MR) method

The moment-ratio matching method uses the ratio of skewness to excess kurtosis to select the gamma shape parameter. Here, excess kurtosis is kurtosis measured relative to the normal-distribution reference value of 3. Let 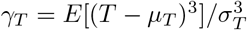 and 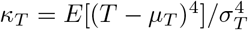 denote the skewness and kurtosis of *T* . The MR-based shape parameter is

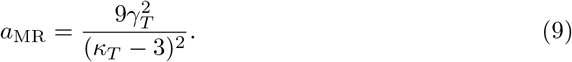

The moments in Eq (9) can be estimated by sampling from the null distribution of *T* through Eqs (4) and (5). The resulting empirical distribution of *T* is then used to estimate its mean *µ*_*T*_, variance 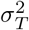, skewness *γ*_*T*_, and kurtosis *κ*_*T*_ for the MR approximation. The estimated mean and variance are then applied to Eq (7).

### Hybrid (HYB) approximation

The hybrid approximation combines MR with the quadratic approximation (Q-approximation) of Zhang and Wu [20]. The Q-approximation adopts a weighted sum of squared normal variables as a surrogate for the distribution of *T* .

By substituting *γ*_*T*_ and *κ*_*T*_ with the Gaussian quadratic form of skewness *γ*_*Q*_ and kurtosis *κ*_*Q*_, the resulting shape parameter is simplified to

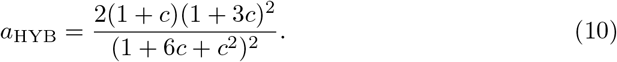

In this work, we implemented HYB using the `R` package `GFisher`, version 0.2.0.

### The adjusted meta-analysis method

Another approach to addressing the sample overlap issue is based on effect sizes. Lin and Sullivan [4] proposed the adjusted meta-analysis method that combines study-specific log-odds estimates 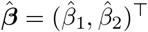, assuming a common effect size. The combined estimate and its variance are

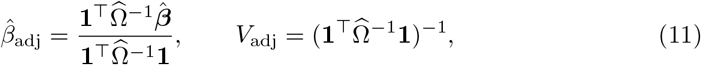

where

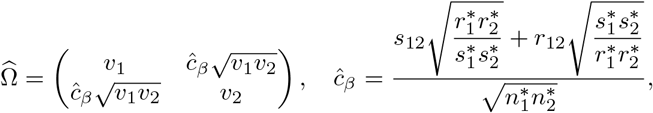

*r*_12_ and *s*_12_ are the number of shared cases and controls, respectively, and 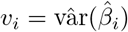for *i* = 1 and 2.

Under the null of no association,

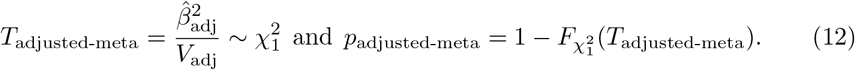

### Mega-analysis as a benchmark

Mega-analysis pools the individual-level data and includes each shared participant only once. For the pooled sample, let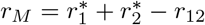and 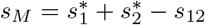 denote the numbers of cases and controls. The score statistic is

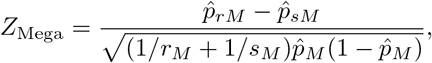

where 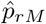 and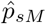are the frequencies of genotype *AA* among pooled cases and controls, and 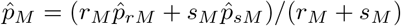). Under the null hypothesis, *Z*_Mega_ is asymptotically standard normal with *p*-value *p*_Mega_ = 2Φ( *Z*_Mega_ ). Mega-analysis based on unique participants serves as the benchmark throughout our analysis.

## Results

### Type I error rates of all tests remain well controlled under modest sample overlap at significance level *α* = 0.05

#### Simulation setup

We first examined the accuracy and power of all the tests using simulated data. The minor allele frequency (MAF) takes value in 0.1, 0.2, 0.3, 0.4, and genotypes were sampled under Hardy–Weinberg equilibrium. We assumed a binary phenotype outcome *Y* and a recessive disease model where *f*_2_ = *P* (*Y* = 1|*AA*) is the disease penetrance probability for genotype *AA* and *f*_1_ = *f*_0_ = *P* (*Y* = 1 *Aa* or *aa*) denotes the disease penetrance probability for genotypes *Aa* and *aa*. The disease prevalence is *K* = *f*_1_(1 − *p*^2^) + *f*_2_*p*^2^. We assumed *K* = 0.1 in our simulation. The probabilities of genotype *AA* among cases and controls are

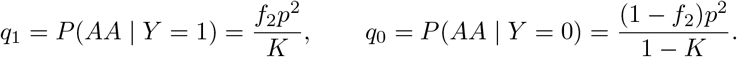

The corresponding probabilities of the combined *Aa/aa* group are 1 −*q*_1_ and 1 −*q*_0_, respectively.

We assumed that there are two overlapping studies each with sample size 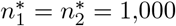and equal number of cases and controls in each study, that is, 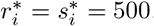. The overlapping sample size *n*_12_ ranges from 50 to 950 participants, corresponding to 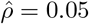 to 0.95. We assume that each sample in Study 1 or Study 2 is equally likely to be included among the overlapping sample (see Section B in S1 File for a detailed description of how the shared and study-specific samples were simulated for predetermined sample sizes). The empirical type 1 error rate and power are evaluated using 10,000 replicates, and evaluated using a nominal threshold *α* = 0.05.

#### Empirical type I error rates

Across all the examined MAFs, GB, HYB, and MR maintained well-controlled empirical T1E rates (Fig 1). The mega-analysis, which serves as the reference analysis using individual-level data, also maintained T1E close to the nominal level. For the classical Fisher’s meta-analysis, T1E remained well controlled when the number of overlapping samples was below 250 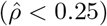. As the degree of sample overlap increased, however, Fisher’s method exhibited progressively inflated T1E, reaching approximately 0.09 when the number of overlapping samples increased to 950 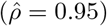. The T1E rate of the adjusted-meta is conservative when MAF = 0.1, which may be related to the assumed recessive disease model. We further examined the accuracy of all tests at more stringent significance levels *α* = 0.01, 0.005 and 0.001 with MAF = 0.3 (Table D in S1 File). The results were consistent with our earlier findings: Fisher exhibited inflated T1E when the sample overlap is greater than 300, while all the other methods maintained controlled T1E.

**Fig 1.**
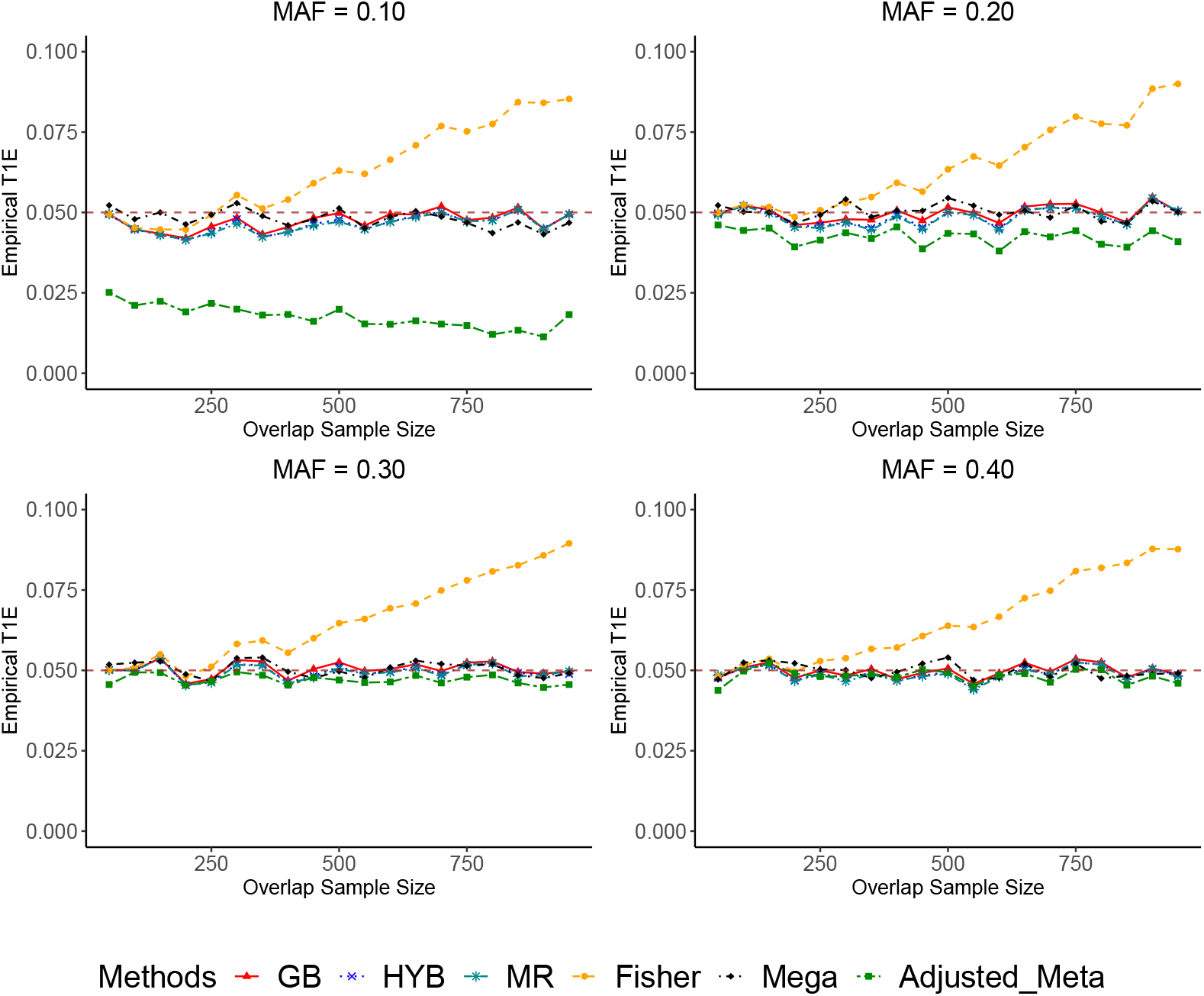
Empirical type I error rates of GB (red), HYB (blue), MR (dark cyan), Fisher (orange), adjusted-meta (green), and mega-analysis (black). Each study includes 1,000 participants, with sample overlap ranging from 50 to 950. The minor allele frequency takes values in {0.1, 0.2, 0.3, 0.4} and genotypes are simulated under Hardy–Weinberg equilibrium. The empirical T1E rates are estimated from 10,000 replicates for each simulation scenario at the significance level of *α* = 0.05. Rejection rates for adjusted-meta are calculated among replicates with defined tests; all other methods use all replicates.

#### Statistical power

We next examined power under the alternative hypothesis with *f*_2_ ∈ {0.12, 0.135, 0.15} . In the power comparison, we also considered a corrected version of the classical Fisher’s method, denoted as “Fisher (corrected)”, that uses the 95th percentile of its simulated null distribution at each overlap size as the rejection threshold. Fig 2 visualises the relative power of all tests evaluated at a significance level *α* = 0.05 using the power of the mega-analysis as a reference.

**Fig 2.**
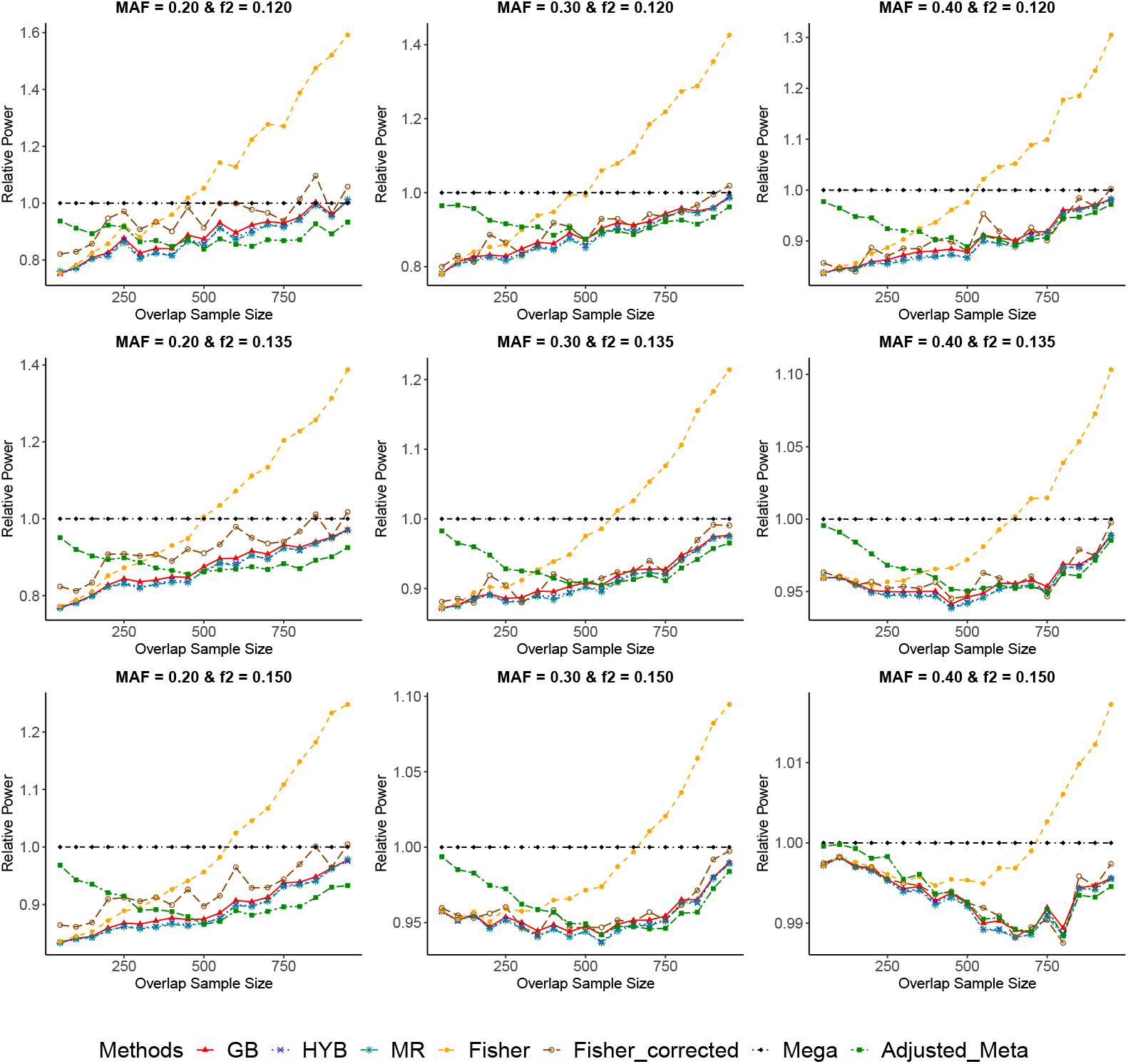
Empirical power relative to mega-analysis of GB, HYB, MR, Fisher, adjusted-meta, and Fisher (corrected). Each study includes 1,000 participants, with sample overlap ranging from 50 to 950. The minor allele frequency takes values in {0.2, 0.3, 0.4}. The disease prevalence is set to *K* = 0.1, and the genotype-specific penetrance *f*_2_ takes values in {0.12, 0.135, 0.15} . The empirical power is estimated from 10,000 replicates for each simulation scenario at a nominal significance level of *α* = 0.05. Relative power is calculated by scaling the empirical power within each simulation scenario to that of the mega-analysis using independent individual-level data.

GB, HYB, and MR showed similar power across the simulation settings. At low to moderate levels of overlap, the three GFisher-based methods generally had lower power than adjusted-meta. This difference narrowed as the sample overlap increases to 500 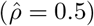and reversed at higher levels of overlap.

The classical Fisher’s method appeared to have the highest power across many settings, but this apparent advantage is largely due to its inflated T1E as shown in Fig 1. Fisher (corrected) exhibited relatively high power across the examined settings while accounting for its T1E inflation. However, the correction relies on an empirical null distribution specific to allele frequency, sample size, and overlap size, making it impractical for large-scale applications such as GWAS.

### Application studies motivated by the HostSeq databank

The HostSeq databank collected whole-genome sequencing and clinical data from 10,059 Canadians with SARS-CoV-2 infection between March 2020 and March 2023 [3]. Several pairs of contributing cohorts shared participants, with the largest overlap observed between BQC19 and IPCO. We used HostSeq genetic data and complementary simulation studies to evaluate five meta-analysis methods, including a chromosome-wide association analysis of BQC19 and IPCO. Mega-analysis based on unique participants served as the benchmark.

#### Re-analysis of six variants from HostSeq with simulated sample overlap

We first analyzed six variants from HostSeq with simulated sample overlap to validate if our findings in the simulation studies hold in real genetic data. The selected loci spanned a range of effect sizes and minor allele frequencies (see Table E in S1 File for details). Three SNPs (rs17763742, rs2496646, rs2834164) had previously been reported to be associated with COVID-19 severity in the HostSeq study [3] or by the Host Genetics Initiative (HGI) [22], while the remaining three variants (rs9990678, rs12345917, rs11064350) showed little evidence of association with the phenotype.

Following the earlier simulation design, we considered two partially overlapping studies of equal sample size, with target overlap rates ranging from 0.1 to 0.9. An 80% subsample of the original data was drawn for each simulation replicate. After rounding the sample sizes, the realized overlap rates 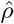 ranged from approximately 0.18 to 0.95. The sample sizes of both studies, shared participant counts, and corresponding 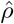 values are provided in Table F in S1 File. Genotypes were extracted using bcftools (v1.21), and participants with missing genotype data were excluded. We assumed a recessive disease model. The procedure was repeated 1,000 times for each combination of locus and overlap level, and the mean *p*-value for each method was reported in Fig 3.

**Fig 3.**
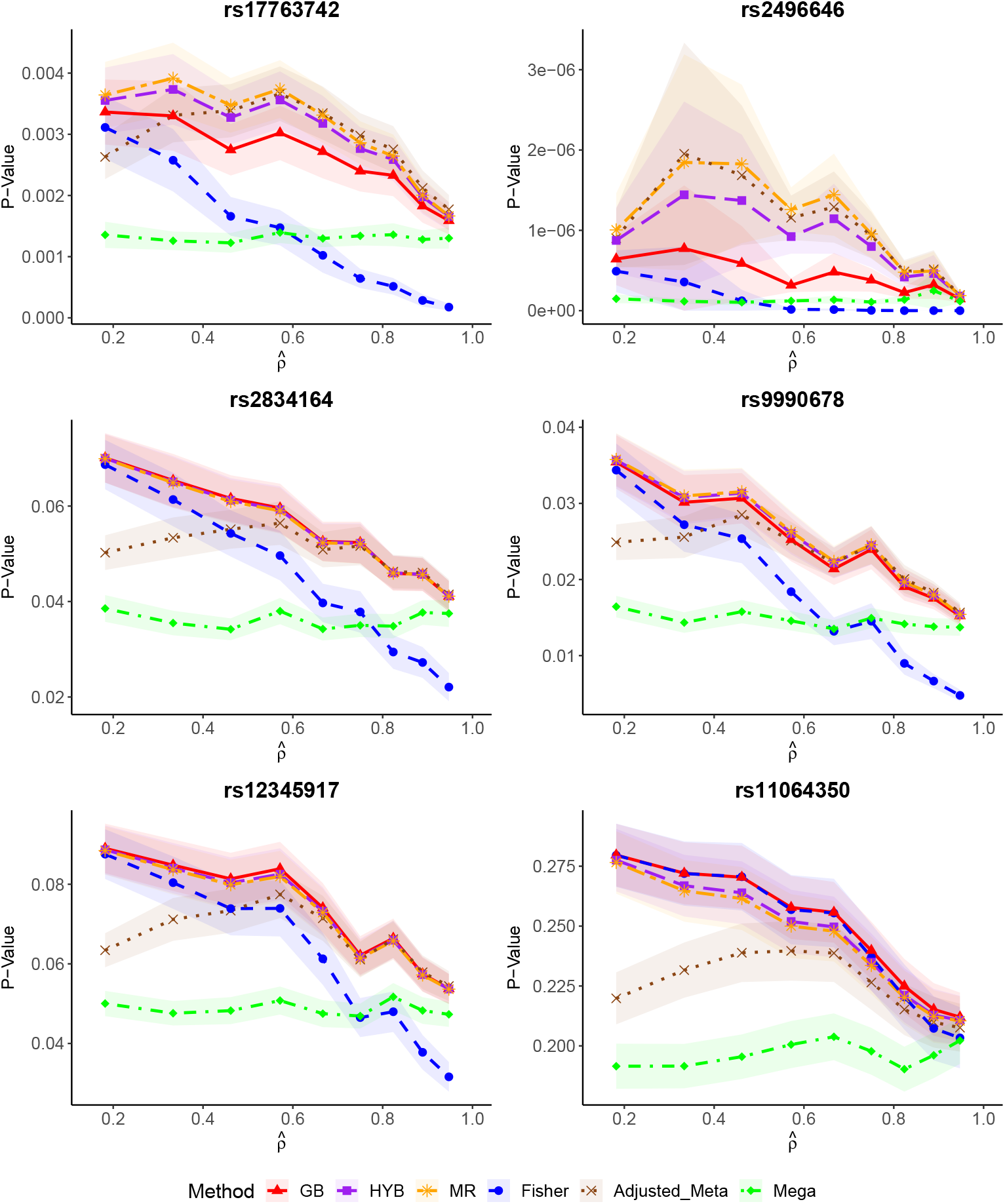
Mean *p*-values across six HostSeq variants with simulated sample overlap. The six methods examined are Fisher, GB, HYB, MR, adjusted-meta, and mega-analysis. For each simulation replicate, 80% of the available samples were randomly selected to construct two partially overlapping studies of equal sample size. The mean *p*-values were calculated from 1,000 replicates for each variant and level of sample overlap. Shaded bands represent the mean *p*-values ±1.96 standard errors, with bounds restricted to [0, 1].

We used the mega-analysis results as the benchmark and assessed the other methods’ agreement with them. Across all six variants examined, mega-analysis (green) exhibited stable *p*-values across different levels of sample overlap. Its small variation observed across overlap levels is due to the subsampling procedure, which resulted in slightly different subsets of unique participants being analyzed in each replicate and at each overlap level. Our mega-analysis results may differ from those reported by HostSeq or HGI (Table E in S1 File) because we assumed a recessive disease model, whereas those analyses assumed an additive model. Nonetheless, the overall strength of the association signals was similar.

Across the six variants, all four methods that account for sample overlap yielded larger *p*-values than the mega-analysis benchmark. This suggests that these methods may be conservative, although the degree of conservativeness diminished as sample overlap increased. Fisher’s method, on the other hand, produced larger *p*-values than the mega-analysis benchmark at low levels of overlap but smaller *p*-values as overlap increased. One exception was observed for the last null variant (rs11064350), where Fisher’s method showed a pattern similar to the three GFisher-based methods, yielding bigger *p*-values than the benchmark across all overlap levels. These results from the real genetic data analysis with simulated sample overlap supported our findings in the earlier simulation studies.

#### A chromosome-wide association study of two HostSeq cohorts with known sample overlap

Our study was motivated by the sample overlap identified among HostSeq cohorts, where Garg et al. [3] identified and removed overlapping individuals. To evaluate the methods using real meta-analysis sample overlap and real genetic data, we conducted a chromosome-wide association analysis (CWAS) of COVID-19 severity using chromosome 21 data from BQC19 and IPCO, the two HostSeq cohorts with the largest number of overlapping participants. BQC19 included 4,001 participants (2,513 cases and 1,488 controls), while IPCO included 183 participants (32 cases and 151 controls). The two cohorts shared 11 participants (8 cases and 3 controls), yielding an estimated overlap correlation of 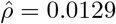 using Eq (3). Because the overlapping individuals were identified, we were able to construct a reference mega-analysis using only unique participants. Following the quality control procedures described in Garg et al. [3] (see Section C in S1 File for details), 674 variants remained for analysis.

We first conducted association analyses separately in BQC19 and IPCO using logistic regression under a recessive disease model, adjusting for the top seven genetic principal components. We used the score tests to obtain the *p*-values. Fisher’s method, GB, HYB, and MR combined transformations of the two *p*-values, with the latter three methods accounting for their correlation. For MR, the moments were estimated empirically using 10^6^ samples drawn from the bivariate normal distribution specified in Eq (5). The adjusted-meta approach combined the effect size estimates obtained from the same logistic regression models and used their corresponding standard errors for variance estimation. The reference mega-analysis adopted the same logistic regression model on the pooled individual-level data with duplicate participants included only once.

Fig 4 compares the *p*-values on −log_10_-scale from MR, adjusted-meta, and Fisher’s method against mega-analysis as the benchmark. MR was selected to represent the three GFisher-based methods because it achieved the closest-to-nominal T1E among them in most of our simulation studies. Adjusted-meta could not test for the association of 71 out of the total 674 variants because the recessive genotype count in case or control was zero. On the remaining 603 variants, all three methods yielded *p*-values similar to those from mega-analysis, with adjusted-meta showing the closest agreement.

**Fig 4.**
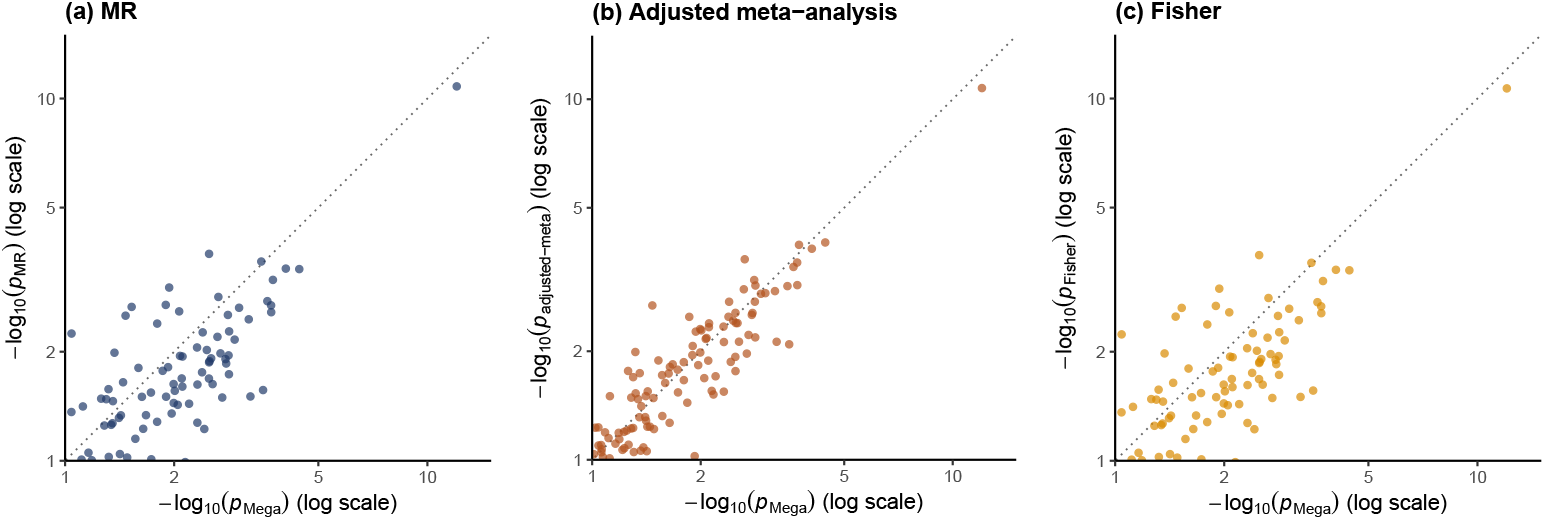
PP-plots comparing chromosome 21 CWAS results from (a) MR, (b) adjusted-meta, and (c) Fisher’s method with mega-analysis as the benchmark. The analysis included 603 variants from the BQC19 and IPCO cohorts for which all methods yielded defined test statistics. The *p*-values are displayed on the −log_10_ scale, with *p*-values *>* 0.1 omitted for better visualisation. The complete PP-plots including all 603 variants are provided in Fig B in S1 File.

Table 2 summarizes the number of chromosome 21 variants with discordant association decisions between the meta-analysis methods and the benchmark mega-analysis across different significance levels. The comparisons were restricted to the 603 variants for which all methods were defined. Fisher’s method and the three GFisher-based methods (GB, MR, and HYB) identified the same significant variants across all examined thresholds. At both the chromosome-wide (*α* = 10^−5^) and genome-wide (*α* = 5 × 10^−8^) significance thresholds, all meta-analysis methods identified the same single significant variant as the benchmark mega-analysis.

**Table 2.** Numbers of variants with discordant significance decisions between the meta-analysis methods and the benchmark mega-analysis in the chromosome 21 CWAS. At each significance threshold *α*, “Significant only by method” reports the number of variants identified as significantly associated by the indicated method but not by mega-analysis, whereas “Significant only by mega-analysis” reports the number of variants identified as significantly associated by mega-analysis but not by the indicated method. Fisher’s method, GB, MR, and HYB identified the same significantly associated variants at each threshold and are therefore grouped together. All comparisons use the 603 variants for which all methods were defined. The Significance level *α* Significant only by method Significant only by mega-analysis Bonferroni-corrected threshold (0.05*/*603 ≈ 8.29 × 10^−5^) is based on the full set of 603 variants.

| Significance level $\alpha$ | Significant only by method | Significant only by mega-analysis |
| --- | --- | --- |
| **Fisher, GB, MR, HYB** |  |  |
| $\alpha = 5 \times 10^{-2}$ | 18 | 26 |
| $\alpha = 5 \times 10^{-3}$ | 8 | 23 |
| $\alpha = 5 \times 10^{-4}$ | 2 | 5 |
| $\alpha = 0.05/603$ | 0 | 1 |
| $\alpha = 1 \times 10^{-5}$ | 0 | 0 |
| $\alpha = 5 \times 10^{-8}$ | 0 | 0 |
| **Adjusted-meta** |  |  |
| $\alpha = 5 \times 10^{-2}$ | 9 | 13 |
| $\alpha = 5 \times 10^{-3}$ | 5 | 11 |
| $\alpha = 5 \times 10^{-4}$ | 1 | 3 |
| $\alpha = 0.05/603$ | 0 | 1 |
| $\alpha = 1 \times 10^{-5}$ | 0 | 0 |
| $\alpha = 5 \times 10^{-8}$ | 0 | 0 |

#### Evaluating test accuracy at genome-wide significance level *α* = 5 × 10^−8^ using the observed BQC19 and IPCO sample overlap

The limited number of variants in the CWAS on chromosome 21 prevented a reliable examination of the test accuracy at the genome-wide significance level *α* = 5 × 10^−8^. We therefore conducted an additional simulation study using the sample sizes and overlap sizes observed in the BQC19 and IPCO cohorts. We simulated Study 1 with 2,513 cases and 1,488 controls, matching BQC19, and Study 2 with 32 cases and 151 controls, matching IPCO. Study 1 and 2 shared 11 participants, matching the observed overlap, but the overlapping individuals were randomly selected without fixing their original case-control composition.

We generated the genotypes that are independent of each individual’s COVID-19 severity status, assuming a minor allele frequency of 0.3 and Hardy–Weinberg equilibrium. We repeated the simulation procedure 10^10^ times to estimate the empirical T1E rates of all methods at the genome-wide significance level *α* = 5 × 10^−8^.

By design, the correlation between the study-specific score test statistics was approximated by 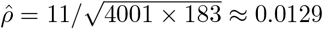 as in Eq (3). The moments of MR were estimated using 5 × 10^7^ random samples from the bivariate normal distribution specified in Eq (5). We also reported the *p*-values based on score tests for both studies. As in the CWAS analysis, the adjusted-meta test statistic was undefined when any genotype count was zero in cases or controls. We therefore calculated its empirical T1E rate using only the replicates for which the test statistic was defined.

Table 3 summarizes the empirical T1E rates of all methods, including the study-specific score tests. Mega-analysis and Study 1 maintained well-controlled T1E at both *α* = 10^−5^ and *α* = 5 × 10^−8^, while Study 2 exhibited substantial T1E inflation at both thresholds. This inflation may be attributed to the small sample size of Study 2 (32 cases and 151 controls). Under the recessive disease model and a MAF of 0.3, the genotype counts in Study 2 are very likely to be sparse, and the asymptotic null distribution of the score test may be inaccurate, particularly in the extreme tail.

**Table 3.** Empirical type I error rates under the observed BQC19 and IPCO sample sizes and sample overlap. Study 1 includes 2,513 cases and 1,488 controls, while Study 2 includes 32 cases and 151 controls. The two studies share 11 participants, randomly selected without fixing their case-control composition. Genotypes were simulated under Hardy–Weinberg equilibrium with a MAF of 0.3. GB, HYB, and MR corrects for sample dependence using 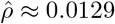 as defined in (3). The empirical T1E rates were estimated from 10^10^ null replicates at significance levels *α* = 10^−5^ and *α* = 5 × 10^−8^. Study-specific score tests were also reported. For adjusted-meta, approximately 4.89% of its test statistics were undefined due to zero genotype counts, and its empirical T1E rates were calculated using only replicates with defined tests. The numbers of rejected tests and corresponding 95% confidence intervals for the empirical T1E rates are provided in Table C in S1 File.

| Method | $\alpha = 10^{-5}$ | $\alpha = 5 \times 10^{-8}$ |
| --- | --- | --- |
| Fisher | $3.502 \times 10^{-5}$ | $6.655 \times 10^{-7}$ |
| GB | $3.497 \times 10^{-5}$ | $6.647 \times 10^{-7}$ |
| HYB | $3.492 \times 10^{-5}$ | $6.624 \times 10^{-7}$ |
| MR | $3.479 \times 10^{-5}$ | $6.592 \times 10^{-7}$ |
| Adjusted-meta | $7.851 \times 10^{-6}$ | $2.912 \times 10^{-8}$ |
| Mega-analysis | $9.738 \times 10^{-6}$ | $4.930 \times 10^{-8}$ |
| Study 1 score test | $9.801 \times 10^{-6}$ | $4.920 \times 10^{-8}$ |
| Study 2 score test | $6.786 \times 10^{-5}$ | $1.526 \times 10^{-6}$ |

The inflated T1E in Study 2 also affected the combined tests. Fisher’s method and all three GFisher-based methods exhibited inflated T1E, with more substantial inflation at *α* = 5 × 10^−8^ than at *α* = 10^−5^. The well-controlled T1E in Study 1 may have partially mitigated the inflation in the combined tests. Adjusted-meta also exhibited inflated T1E, although to a lesser extent. This difference may be partly explained by its exclusion of simulation replicates with zero genotype counts, which represented the sparse genotype scenarios where the other methods may be more prone to T1E inflation.

#### Evaluating test accuracy at stringent significance thresholds in GWAS-sized simulation studies

In the simulation studies based on the observed sample sizes and overlap between BQC19 and IPCO, the sparse genotype counts (due to the small sample size of IPCO) might have caused the inflated T1E at *α* = 10^−5^ and *α* = 5 × 10^−8^ for all the meta-analysis methods. To minimize the impact of genotype sparsity, we conducted additional simulations with larger sample sizes to evaluate T1E at more stringent significance thresholds. We considered two larger studies each with 20,000 participants, and overlap sizes of 2,000 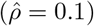 and 6,000 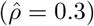. Each study included equal numbers of cases and controls. We simulated genotypes independent of the binary phenotype outcome, assuming Hardy–Weinberg equilibrium and MAFs of 0.1, 0.2, 0.3, 0.4, and 0.5. The detailed data generation procedure is described in Section B in S1 File. Empirical T1E rates were estimated using 10^10^ independent replicates for nominal significance thresholds *α* = 10^−5^ and *α* = 5 × 10^−8^. We compared all five meta-analysis methods and mega-analysis as the benchmark. All methods were implemented as in the earlier simulations based on the BQC19 and IPCO sample size and overlap size.

Table 4 summarizes the empirical T1E rates of all methods estimated at *α* = 10^−5^ and *α* = 5 × 10^−8^. Fig A in S1 File visualizes the corresponding empirical-to-nominal error ratios at MAF 0.3 with 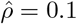 and 0.3. Mega-analysis generally maintained T1E rates close to the nominal levels. At MAF= 0.1, mega-analysis exhibited conservative T1E at both significance thresholds and overlap levels, which may be caused by genotype count imbalance under a recessive design (see Discussion for more details). Fisher’s method exhibited inflated T1E across all scenarios. The inflation became substantial as the sample overlap increased from 2,000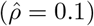) to 6,000 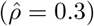 participants. At the smaller overlap level 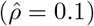, its T1E inflation was similar to that of GB, but this difference between became considerably larger at the higher overlap level 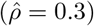.

**Table 4.** Empirical type I error rates evaluated at chromosome-wide significance level *α* = 10^−5^ and genome-wide significance level *α* = 5 × 10^−8^. Each study included 20,000 participants, with equal numbers of cases and controls. The two studies shared 2,000 participants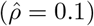 or 6,000 participants 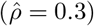. Genotypes were simulated under Hardy–Weinberg equilibrium with MAF = {0.1, 0.2, 0.3, 0.4, 0.5} Empirical T1E were estimated from separate sets of 10^10^ null replicates for each simulation scenario and significance threshold. The number of rejected tests and pointwise 95% intervals are provided in Table B in S1 File.

| $\hat{\rho}$ | MAF | Fisher | GB | HYB | MR | Adjusted-meta | Mega-analysis |
| --- | --- | --- | --- | --- | --- | --- | --- |
| $\alpha = 10^{-5}$ | | | | | | | |
| 0.1 | 0.1 | $1.202 \times 10^{-5}$ | $1.105 \times 10^{-5}$ | $9.890 \times 10^{-6}$ | $9.566 \times 10^{-6}$ | $7.006 \times 10^{-6}$ | $9.331 \times 10^{-6}$ |
| | 0.2 | $1.311 \times 10^{-5}$ | $1.210 \times 10^{-5}$ | $1.086 \times 10^{-5}$ | $1.051 \times 10^{-5}$ | $9.255 \times 10^{-6}$ | $9.810 \times 10^{-6}$ |
| | 0.3 | $1.335 \times 10^{-5}$ | $1.230 \times 10^{-5}$ | $1.107 \times 10^{-5}$ | $1.070 \times 10^{-5}$ | $9.735 \times 10^{-6}$ | $9.973 \times 10^{-6}$ |
| | 0.4 | $1.334 \times 10^{-5}$ | $1.231 \times 10^{-5}$ | $1.106 \times 10^{-5}$ | $1.070 \times 10^{-5}$ | $9.879 \times 10^{-6}$ | $1.001 \times 10^{-5}$ |
| | 0.5 | $1.349 \times 10^{-5}$ | $1.244 \times 10^{-5}$ | $1.118 \times 10^{-5}$ | $1.083 \times 10^{-5}$ | $9.954 \times 10^{-6}$ | $9.999 \times 10^{-6}$ |
| 0.3 | 0.1 | $4.165 \times 10^{-5}$ | $2.232 \times 10^{-5}$ | $1.197 \times 10^{-5}$ | $9.289 \times 10^{-6}$ | $6.525 \times 10^{-6}$ | $9.204 \times 10^{-6}$ |
| | 0.2 | $4.464 \times 10^{-5}$ | $2.413 \times 10^{-5}$ | $1.304 \times 10^{-5}$ | $1.014 \times 10^{-5}$ | $9.043 \times 10^{-6}$ | $9.806 \times 10^{-6}$ |
| | 0.3 | $4.533 \times 10^{-5}$ | $2.452 \times 10^{-5}$ | $1.334 \times 10^{-5}$ | $1.040 \times 10^{-5}$ | $9.641 \times 10^{-6}$ | $9.879 \times 10^{-6}$ |
| | 0.4 | $4.558 \times 10^{-5}$ | $2.466 \times 10^{-5}$ | $1.338 \times 10^{-5}$ | $1.044 \times 10^{-5}$ | $9.824 \times 10^{-6}$ | $9.939 \times 10^{-6}$ |
| | 0.5 | $4.547 \times 10^{-5}$ | $2.471 \times 10^{-5}$ | $1.345 \times 10^{-5}$ | $1.048 \times 10^{-5}$ | $9.901 \times 10^{-6}$ | $9.949 \times 10^{-6}$ |
| $\alpha = 5 \times 10^{-8}$ | | | | | | | |
| 0.1 | 0.1 | $6.890 \times 10^{-8}$ | $5.830 \times 10^{-8}$ | $4.900 \times 10^{-8}$ | $4.630 \times 10^{-8}$ | $2.110 \times 10^{-8}$ | $4.040 \times 10^{-8}$ |
| | 0.2 | $8.840 \times 10^{-8}$ | $7.840 \times 10^{-8}$ | $6.640 \times 10^{-8}$ | $6.260 \times 10^{-8}$ | $4.440 \times 10^{-8}$ | $5.200 \times 10^{-8}$ |
| | 0.3 | $8.630 \times 10^{-8}$ | $7.620 \times 10^{-8}$ | $6.330 \times 10^{-8}$ | $6.090 \times 10^{-8}$ | $4.040 \times 10^{-8}$ | $4.570 \times 10^{-8}$ |
| | 0.4 | $9.340 \times 10^{-8}$ | $8.170 \times 10^{-8}$ | $6.900 \times 10^{-8}$ | $6.670 \times 10^{-8}$ | $5.100 \times 10^{-8}$ | $5.190 \times 10^{-8}$ |
| | 0.5 | $9.290 \times 10^{-8}$ | $8.200 \times 10^{-8}$ | $6.790 \times 10^{-8}$ | $6.450 \times 10^{-8}$ | $4.970 \times 10^{-8}$ | $5.260 \times 10^{-8}$ |
| 0.3 | 0.1 | $5.482 \times 10^{-7}$ | $2.052 \times 10^{-7}$ | $6.680 \times 10^{-8}$ | $4.300 \times 10^{-8}$ | $1.930 \times 10^{-8}$ | $3.760 \times 10^{-8}$ |
| | 0.2 | $6.433 \times 10^{-7}$ | $2.407 \times 10^{-7}$ | $8.760 \times 10^{-8}$ | $5.740 \times 10^{-8}$ | $4.220 \times 10^{-8}$ | $5.000 \times 10^{-8}$ |
| | 0.3 | $6.754 \times 10^{-7}$ | $2.533 \times 10^{-7}$ | $8.920 \times 10^{-8}$ | $5.680 \times 10^{-8}$ | $4.560 \times 10^{-8}$ | $4.980 \times 10^{-8}$ |
| | 0.4 | $6.710 \times 10^{-7}$ | $2.592 \times 10^{-7}$ | $9.100 \times 10^{-8}$ | $5.950 \times 10^{-8}$ | $4.920 \times 10^{-8}$ | $4.500 \times 10^{-8}$ |
| | 0.5 | $6.678 \times 10^{-7}$ | $2.552 \times 10^{-7}$ | $9.280 \times 10^{-8}$ | $6.220 \times 10^{-8}$ | $5.250 \times 10^{-8}$ | $5.330 \times 10^{-8}$ |

Among the three GFisher-based methods, GB consistently exhibited the greatest T1E inflation. HYB substantially reduced this inflation relative to GB, while MR achieved the closest T1E rates to the nominal levels. At MAF = 0.1, MR exhibited deflated T1E, consistent with the T1E pattern observed in mega-analysis, but to a lesser extent. Adjusted-meta exhibited conservative T1E rates across most scenarios, particularly at lower MAFs.

Each study includes 20,000 participants, with equal numbers of cases and controls. The two studies share 2,000 participants 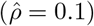 or 6,000 participants 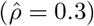. Empirical T1E rates were estimated from separate sets of 10^10^ null replicates for each simulation scenario and significance threshold. The results at *α* = 5 × 10^−8^ were obtained from the initial simulations, while those at *α* = 10^−5^ used newly simulated genotypes with the same previously fitted distributions. Fisher’s method, GB, HYB, and MR combine study-specific score-test *p*-values, while adjusted-meta uses Wald estimates of log-odds ratios. Mega-analysis applies a score test to the pooled data with unique participants. All methods were defined in every replicate. The numbers of rejections and corresponding pointwise 95% confidence intervals are provided in Table B in S1 File.

## Discussion

Motivated by the sample overlap observed in HostSeq cohorts, we investigated how sample overlap in GWAS meta-analysis affect the results and evaluated existing methods that account for sample dependence. We compared the classical Fisher’s method and four methods that account for sample overlap, using mega-analysis based on unique participants as the benchmark. Through extensive simulation studies and applications to HostSeq data, we demonstrated that the impact of sample overlap on association results depends on the degree of overlap, sample sizes of the contributing studies, and genotype frequencies of the variants of interest.

The classical Fisher’s method controlled T1E at *α* = 0.05 when no more than 250 participants were shared between studies each with 1,000 participants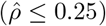. In the analyses using real HostSeq genetic data with simulated sample overlap, Fisher’s method also yielded concordant *p*-values with mega-analysis, even with 2,000 overlapping participants between studies of ∼4,400 participants each 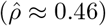. In the CWAS of BQC19 and IPCO that shared 11 participants, Fisher’s method and mega-analysis identified the same significant variants (among the 674 analyzed variants) at the Bonferroni-corrected, chromosome-wide (*α* = 10^−5^), and genome-wide (*α* = 5 × 10^−8^) significance thresholds. However, in the GWAS-sized simulations using 10^10^ replicates, Fisher’s method exhibited inflated T1E at both thresholds, even with 2,000 overlapping participants between studies of 20,000 participants each 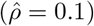

The three GFisher-based methods, generalized Brown’s method, moment-ratio matching method and the hybrid method, generally exhibited similar performance, except in the GWAS-sized simulations at stringent significance thresholds. All three methods maintained well-controlled T1E rates and achieved comparable power, both evaluated at *α* = 0.05. In the analysis of six HostSeq loci with simulated sample overlap, all three methods yielded similar mean *p*-values, consistently larger than those from mega-analysis, although the differences diminished as the overlap size increased. In the GWAS-sized simulation study, MR achieved the closest empirical T1E to the nominal level *α* = 5 × 10^−8^ among the three methods, although some inflation remained.

The adjusted-meta approach exhibited conservative T1E at MAF = 0.1 evaluated using a significance level of *α* = 0.05. In the analysis of six loci with simulated overlap, adjusted-meta yielded mean *p*-values closer to those of mega-analysis than the three GFisher-based methods. In the GWAS-sized simulations at stringent significance thresholds, adjusted-meta exhibited conservative T1E at lower MAFs, but its empirical T1E approached the nominal levels at MAFs of 0.4 and 0.5 under both chromosome-wide and genome-wide significance thresholds.

When adjustment for dependence due to sample overlap is desired, we recommend MR or adjusted-meta, although both methods have limitations. At the genome-wide significance threshold (*α* = 5 × 10^−8^), MR exhibited modest T1E inflation when MAF≥ 0.2, while adjusted-meta tended to be conservative, especially at lower MAFs. MR relies on moment estimation based on the empirical distribution of *T*, and a sufficiently large number of random samples is needed to ensure accurate moment estimation. On the other hand, adjusted-meta is undefined when any genotype count is zero and requires information on genotype counts in case and control groups among overlapping individuals.

An important practical question is how much sample overlap can be tolerated without correction. Fisher’s method maintained well-controlled T1E at *α* = 0.05 with up to 250 overlapping participants between studies each of size 1,000. In the BQC19/IPCO CWAS, the 11 overlapping participants 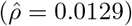 had little impact on association decisions at Bonferroni-corrected, chromosome-wide and genome-wide significance thresholds. Our GWAS-sized simulations showed Fisher’s T1E inflation at *α* = 10^−5^ and *α* = 5 × 10^−8^ with 2,000 overlapping participants between studies of 20,000 participants each. Such substantial overlap may be less representative of incidental duplication between independently recruited cohorts and warrants examining whether the overlap-at-random assumption underlying our correlation approximation is reasonable. In summary, our findings suggest that the need for correction should be assessed based on the number of overlapping participants and individual study sizes. For very small incidental overlaps, Fisher’s method may remain a reasonable practical choice without correction, whereas substantial overlap warrants adjustment for sample dependence.

As with any meta-analysis, the reliability of the meta-analysis results depends on the accuracy of the individual study analyses. The methods examined in this study combine either study-specific *p*-values or effect estimates, both of which may be affected by low genotype counts. Low genotype counts can lead to unstable variance estimates and compromise the asymptotic distributions of the test statistics under the null hypothesis. Under our assumed recessive genetic model, in the simulation studies with MAF= 0.1, the genotype frequency of *AA* is 0.01, resulting in a substantial imbalance between the two genotype groups (*AA* vs. *Aa* or *aa*). Therefore, the results at MAF = 0.1 should be interpreted with caution. Moreover, small genotype counts can arise even at higher MAFs when individual studies have small sample sizes, as in the IPCO-related simulations. In such settings, inaccurate study-specific results may propagate to the combined results, regardless of whether sample overlap is appropriately accounted for.

An important future direction for our work is to adjust for the dependence induced by non-random sample overlap, such as the common practice of reusing controls across GWAS. Such settings require an appropriate estimator of the correlation between study-specific test statistics that accounts for the phenotype outcome of the overlapping participants. Although our analyses here focused on a recessive genetic model, the framework can also be extended to additive and dominant genetic models.

## Supporting information

**S1 File. Supporting information**. Derivation of the correlation between study-level score statistics (Section A); generation of partially overlapping case–control studies (Section B); quality control for the BQC19/IPCO chromosome-wide association analysis (Section C); genotype-count notation (Table A); complete results at two stringent significance thresholds (Table B); complete results for the simulation with 11 shared participants (Table C); empirical type I error rates at four significance levels (Table D); overview of the six selected loci (Table E); study sizes and estimated correlations in the imposed-overlap experiment (Table F); calibration at two significance thresholds (Fig A); and comparison of chromosome 21 association results across all 603 commonly analyzed variants (Fig B).

